# A human monoclonal antibody targeting a conserved pocket in the SARS-CoV-2 receptor-binding domain core

**DOI:** 10.1101/2020.09.30.318261

**Authors:** Juliette Fedry, Daniel L. Hurdiss, Chunyan Wang, Wentao Li, Gonzalo Obal, Ieva Drulyte, Stuart C. Howes, Frank J.M. van Kuppeveld, Friedrich Förster, Berend-Jan Bosch

## Abstract

SARS-CoV-2 has caused a global outbreak of severe respiratory disease (COVID-19), leading to an unprecedented public health crisis. To date, there has been over thirty-three million diagnosed infections, and over one million deaths. No vaccine or targeted therapeutics are currently available. We previously identified a human monoclonal antibody, 47D11, capable of cross-neutralising SARS-CoV-2 and the related 2002/2003 SARS-CoV *in vitro*, and preventing SARS-CoV-2 induced pneumonia in a hamster model. Here we present the structural basis of its neutralization mechanism. We describe cryo-EM structures of trimeric SARS-CoV and SARS-CoV-2 spike ectodomains in complex with the 47D11 Fab. These data reveal that 47D11 binds specifically to the closed conformation of the receptor binding domain, distal to the ACE2 binding site. The CDRL3 stabilises the N343 glycan in an upright conformation, exposing a conserved and mutationally constrained hydrophobic pocket, into which the CDRH3 loop inserts two aromatic residues. Interestingly, 47D11 preferentially selects for the partially open conformation of the SARS-CoV-2 spike, suggesting that it could be used effectively in combination with other antibodies that target the exposed receptor-binding motif. Taken together, these results expose a cryptic site of vulnerability on the SARS-CoV-2 RBD and provide a structural roadmap for the development of 47D11 as a prophylactic or post-exposure therapy for COVID-19.

## Introduction

The severe acute respiratory syndrome coronavirus 2 (SARS-CoV-2) emerged from a zoonotic event in China, late 2019(*1*). To date, the resulting coronavirus induced disease 19 (COVID-19) pandemic has been responsible for over 33 million infections and over a million deaths, as of September 29, 2020(https://covid19.who.int/). SARS-CoV-2 and SARS-CoV, another highly lethal respiratory pathogen which emerged in 2002/2003(*2*), belong to the *Sarbecovirus* subgenus (genus *Betacoronavirus*, family *Coronaviridae*)(*3*). At present, no targeted therapeutics have been approved for COVID-19, meaning there is an urgent clinical need for potent antiviral therapies to halt the spread of SARS-CoV-2 and to pre-empt future outbreaks caused by SARS-like viruses. Antibodies are a promising class of drugs for combatting infectious diseases and have shown therapeutic efficacy for a number of viruses(*4, 5*), including in the treatment of SARS and COVID-19(*6, 7*). Such antibodies function by targeting vulnerable sites on viral surface proteins.

The coronavirus trimeric spike (S) glycoprotein, located on the viral envelope, is the key mediator of viral entry into host cells. The spike protein consists of two main parts: S1 is involved in receptor binding and S2 is the membrane fusion domain. The S1 domain itself is further subdivided into an N-terminal domain (NTD, or S1A) and a receptor binding domain (RBD, or S1B)(*8, 9*). The spike proteins of SARS-CoV-2 (SARS2-S; 1273 residues, strain Wuhan-Hu-1) and SARS-CoV (SARS-S, 1255 residues, strain Urbani) exhibit 77.5% identity in their primary amino acid sequence and are structurally conserved. The spike trimer exists in equilibrium between a closed conformation, where all three RBD are lying flat, and a partially open conformation, where one RBD stands upright and is exposed for receptor engagement(*10–12*). Both viruses use the human angiotensin converting enzyme 2 (ACE2) protein as a host receptor, with binding mediated through interactions with the receptor-binding motif (RBM) located on the RBD, and the N-terminal helix of ACE2(*13*). The spike-mediated fusion of viral and cellular membranes is tightly regulated and triggered by a cascade of preceding events. The first step involves the attachment of SARS-CoV-2 to the target cell surface via the interaction between spike and ACE2(*13, 14*). In the second step, the spike protein needs to be primed for membrane fusion by host proteases (e.g. cellular transmembrane serine protease 2) which cleave the spike at multiple sites(*15*), enabling shedding of S1. Finally, the free S2 catalyses the fusion of the viral and the host membranes(*16, 17*), causing the release of the viral genome into the host cell cytoplasm.

The S glycoprotein is the primary target for neutralising antibodies, making it the main target for vaccine development(*18*). Indeed, a number of SARS-CoV-2 neutralising antibodies have now been described(*19–33*). However, comparatively few cross-neutralising antibodies have been reported(*20, 25, 30, 34*), of which only a handful have been structurally characterised(*29, 35–37*). The most commonly identified antibodies neutralize coronaviruses by binding to the receptor interaction site in S1, blocking receptor interactions and/or promoting premature conformational change of spike to the post-fusion state. However, a smaller number of antibodies have been reported to bind sites which are distal to the ACE2 binding site. Such antibodies target the RBD-core(*27, 35, 37, 38*), or even the NTD(*39*). Several promising vaccines are currently being developed but it is estimated that at least a year will be needed before they can be introduced on the market. Hence there is an urgent need for characterized antibodies to form cocktails for the treatment by passive immunization of COVID-19 patients. Combined structural and functional studies are thus required to determine the epitopes and investigate the molecular mechanisms of SARS-CoV-2 neutralizing antibodies. Moreover, such studies may identify cryptic sites of vulnerability which can guide vaccine and antiviral development(*40*).

We recently reported the first human monoclonal antibody, 47D11, capable of cross-neutralising SARS-CoV and SARS-CoV-2 at 1.3 and 3.8 nM, respectively(41), Moreover, recent pre-clinical studies show that 47D11 protects against lower respiratory tract disease in a hamster model(42). However, its mode of engagement with the spike protein remained unclear. We thus employed structural and functional studies to decipher the molecular basis for 47D11-mediated neutralisation.

## Results

### 47D11 specifically binds to the closed receptor binding domain

To understand how 47D11 binds to the SARS-CoV and SARS-CoV-2 spike proteins, we used cryo-electron microscopy (cryo-EM) to determine structures of prefusion stabilised ectodomain trimers in complex with the 47D11 Fab fragment. The resulting cryo-EM maps have global resolutions of 3.8 Å and 4.0 Å resolution for SARS-S and SARS2-S, respectively (Supplementary Figure 1A-F). For previously reported apo S trimers, both the open and closed conformation are observed, with the latter being predominant (56% for SARS and 67% for SARS2(*11, 43*)). Upon incubation with 47D11, only the closed conformation of the SARS spike was observed, with stochiometric binding of 47D11 to each RBD (Figure 1A). Interestingly, for SARS-CoV-2, only the partially open conformation of spike was observed, with one Fab bound to each of the closed RBDs, and the remaining open RBD unoccupied and, in principle, accessible to ACE2 binding (Figure 1B). The sub stochiometric binding observed for SARS2-S may partially explain our previous observations that 47D11 binds to the SARS-S with higher affinity than SARS2-S (equilibrium dissociation constant [K_D_] of 0.745 nM and 10.8 nM, respectively)(*41*). To understand why 47D11 favours different spike conformations for SARS-CoV and SARS-CoV-2, we first superposed the Fab-bound structures with their apo counterparts. Compared to the apo partially open SARS2-S structure, the RBDs are less compact when 47D11 is bound (Figure 2A). The apo conformation of the closed RBD would preclude binding of 47D11 through steric hindrance. To accommodate the bound Fab, this RBD shifts outwards by ~7 Å (Figure 2B). Unlike S309 and H014(*35, 38*), two other RBD-core targeting SARS neutralizing antibodies, there was no indication from our cryo-EM data that 47D11 can bind to the open conformation of the SARS2-S RBD. In line with this, superimposition of open and closed SARS2-S RBDs revealed that 47D11 would clash with the adjacent N-terminal domain (NTD) and the N331 glycan in the latter conformation (Figure 2C). Similar to SARS2-S, the RBDs of the 47D11 bound SARS-S are also less compact than the reported apo fully closed structure (Figure 2D). However, in contrast to SARS2-S, there is a potential stabilising salt bridge between D463, located on the receptor binding ridge (RBR), and R18 on the 47D11 light chain (Figure 2E). Indeed, the RBR exhibits the most prominent structural differences between SARS2-S and SARS-S(*13*). This epitope distal loop, located within the ACE2-binding region, contains an essential disulfide bridge in both viruses, but is more compact in SARS-S. In order to test whether the epitope distal RBR impacts binding of 47D11 to the SARS-S and SARS2-S, we swapped loop residues 470-490 (SARS2-S numbering) and produced chimeric ectodomains. In support of our hypothesis, the SARS2-S, containing the SARS-S RBR loop, exhibited increased binding to 47D11. However, we did not observe an equivalent loss of binding for the chimeric SARS-S, suggesting that other differences in protein sequence or quaternary structure may be involved (Figure 2F). Taken together, our data shows that 47D11 binding to the closed RBDs of the trimeric spike protein has differing outcomes for SARS-S and SARS2-S, trapping them in the fully closed and the partially open conformation, respectively (Figure S2).

**Figure 1:**
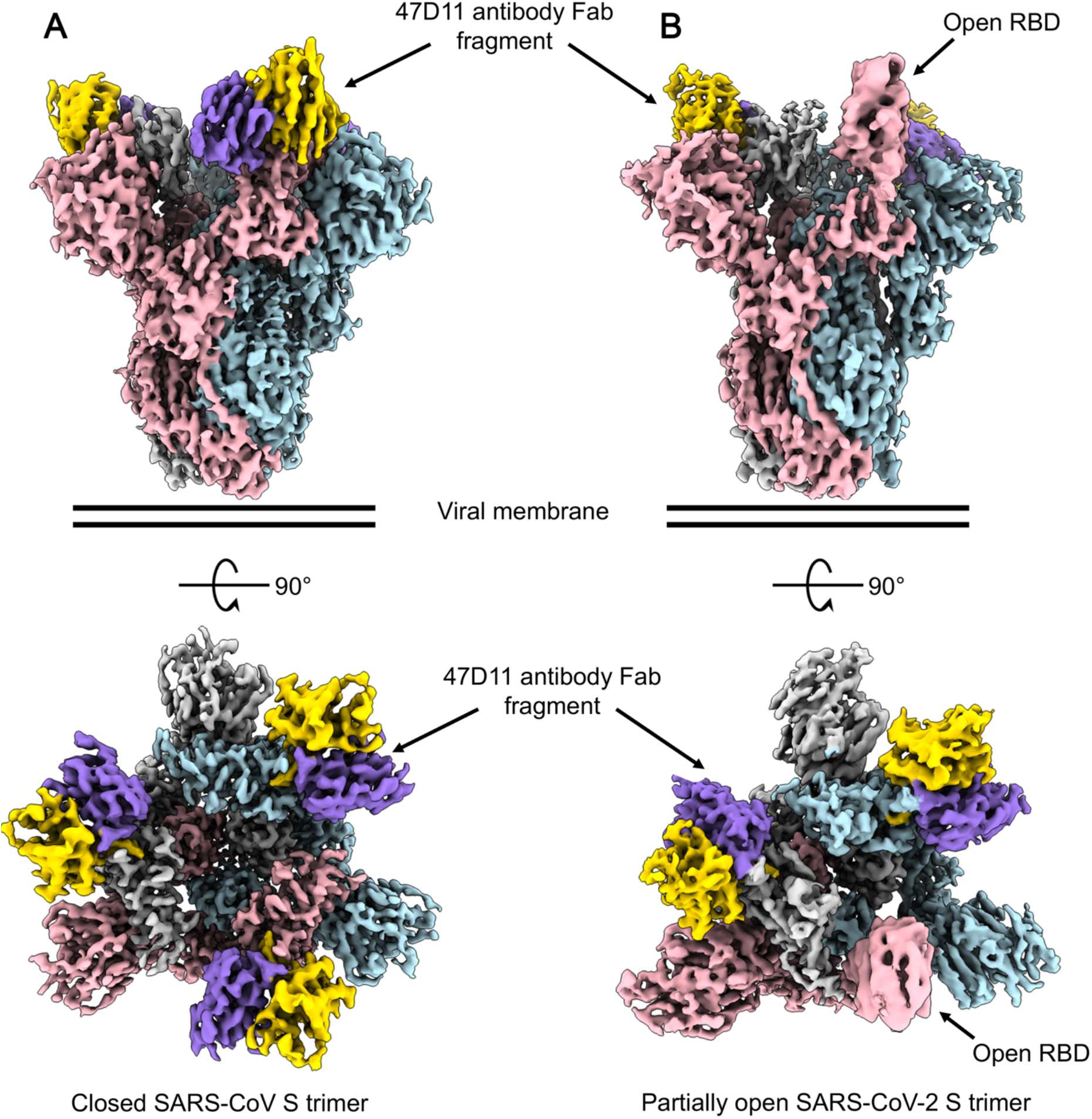
47D11 has differing conformational selectivity for the SARS-CoV and SARS-CoV-2 spike. A) Surface rendering of the fully closed SARS spike bound to three 47D11 antibody Fab fragments, shown as two orthogonal views. (B) Surface rendering of the partially open SARS2 spike in complex with two 47D11 antibody Fab fragments, shown as two orthogonal views. The spike protomers are coloured pink, blue and grey, and the 47D11 heavy and light chain are coloured yellow and purple, respectively. For clarity, only the Fab variable region is shown.

**Figure 2:**
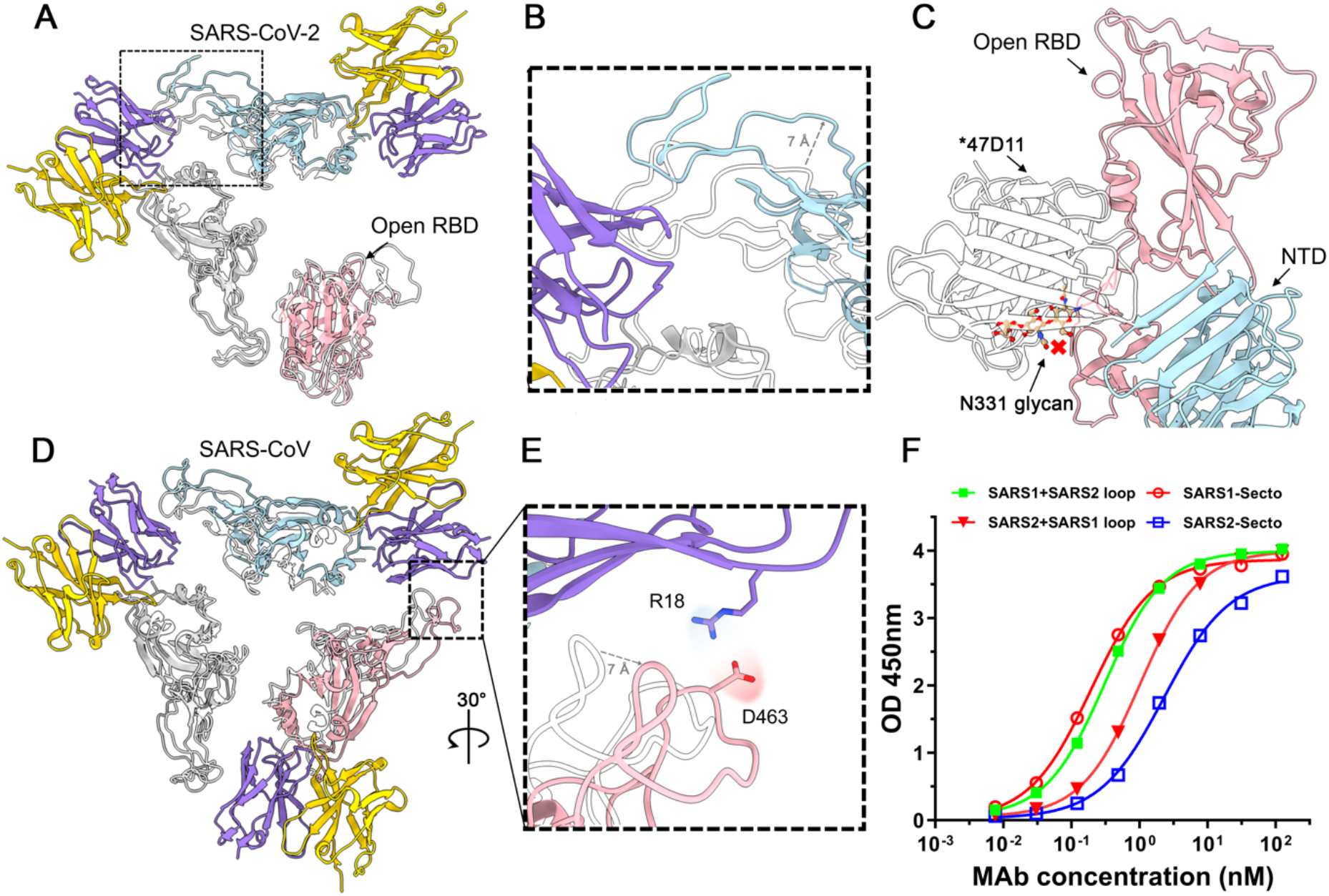
47D11 binds specifically to the closed RBD and prevents their full compaction. A) Top view of the 47D11 bound SARS2 spike. The spike protomers are coloured pink, blue and grey, and the 47D11 heavy and light chain are coloured yellow and purple, respectively. Glycans, and the N-terminal domain, are omitted for clarity and only the Fab variable region is shown. The superposed structure of the partially open apo SARS2 spike (PDB ID: 6ZGG) is shown as a silhouette. B) Zoomed in view of the boxed region in panel A. C) Zoomed in view of the SARS2 open RBD and adjacent NTD. The overlaid 47D11 Fab is shown semitransparent and the N343 glycan is shown in ball-and-stick representation and coloured tan. D) Top view of the 47D11 bound SARS2 spike coloured as shown in panel A. The superposed structure of the closed apo SARS spike (PDB ID: 5XLR) is shown as a silhouette. E) Zoomed in view of the boxed region in panel D, showing a putative salt bridge between the 47D11 variable light chain and the RBD loop. F) ELISA-binding curves of 47D11 binding to wildtype and loop swapped spike ectodomains.

### 47D11 targets a conserved hydrophobic pocket in the RBD

The 47D11 epitope is distinct from the ACE2 binding site (Figure 3A), consistent with our recently reported functional data(*41*). The protein/glycan epitope is located on the core domain of the SARS-S and SARS2-S RBD. As expected, the mode of binding is highly similar for SARS-S and SARS2-S (Figure S3A), with the aligned 47D11:RBD complexes having an RMSD value of 1.4 Å. The paratope is composed of CDRL3 and CDRH3 loops, that form a primarily hydrophobic interaction with the RBD surface of ~830 Å^2^ and ~800 Å^2^ for SARS-S and SARS2-S, respectively. The side chain of 47D11 CDRL3 tryptophan W94 stacks against the N330/N343 (SARS/SARS2) glycan tree, contributing to its stabilization in an upright conformation (Figure 3B). This reveals a hydrophobic pocket into which the CDRH3 loop projects, allowing Fab residues W102 and F103 to interact with RBD core residues F338, F342, Y365, V367, L368, F374 and W436 (F325, F329, Y352, V354, L355, F361 and W423 in SARS-S) – figure 3B and S3B. Interestingly, this pocket is normally shielded by the N343 glycan in previously reported SARS2-S structures (Figure 3C-D)(*11, 12*). In order to accommodate the CDRH3 loop residues, the helix encompassing residues 365-370 is displaced outwards by 2 Å, creating 55 Å^3^ of solvent accessible volume which is not present in the apo RBD (Figure 3C-D and Figure S3C-D). Of note, the region directly below this hydrophobic pocket was recently shown bind to linoleic acid, which stabilises the closed conformation of spike by spanning two adjacent RBDs(*44*). However, the distance between the 47D11-bound RBDs is too great to be bridged by linoleic acid. Consistent with this, no density consistent with linoleic acid was present in any of our reconstructions.

**Figure 3:**
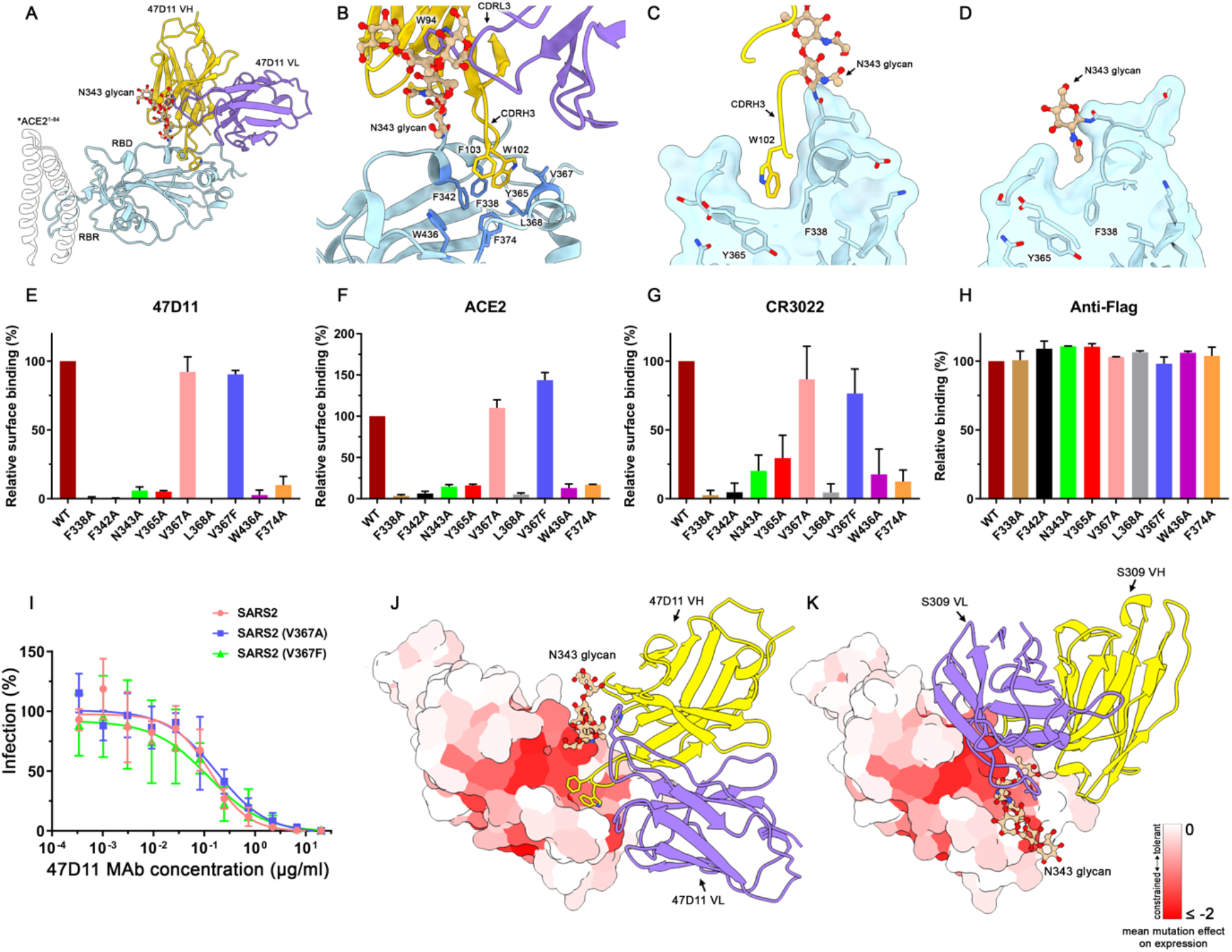
The 47D11 epitope comprises a mutationally constrained hydrophobic pocket which is normally shielded by glycan N343. A) Ribbon diagram of the SARS2-S receptor-binding domain (RBD) in complex with the 47D11 antibody Fab fragment. For comparison, residues 1-84 of the RBD bound ACE2 (PDB ID: 6M0J) are shown as a silhouette. B) Closeup view of the 47D11 epitope with the hydrophobic pocket residues shown as sticks and coloured dark blue. The N343 glycan is shown in ball-and-stick representation and coloured tan. For clarity, only the core pentasaccharide is shown. C) Slice through the surface rendered 47D11 bound SARS2-S RBD. D) Equivalent view as shown in panel C for the apo RBD (PDB ID:6VYB). E) Relative surface binding of 47D11, (F) ACE2, (G) CR3022 and (H) an anti-FLAG antibody to full-length SARS2 spike epitope mutants, determined by fluorescence-activated cell sorting. I) Antibody-mediated neutralization of infection of luciferase-encoding VSV particles pseudotyped with wild-type, V367A or V367F SARS2-S. J) Surface representation of the 47D11 bound SARS2-S RBD coloured according to mean mutation effect on expression (red indicates more constrained)(*46*). The Fab is shown as a ribbon diagram. K) As shown in E for the S309 bound SARS2 RBD.

To verify the 47D11 epitope, we introduced alanine mutations at each of the identified contact residues in the context of full-length spike protein. In addition, a spike mutant with the naturally occurring V367F minority variant was generated(*45*). Binding of 47D11 to surface expressed wildtype and mutant spike proteins was assessed by flow cytometry. Soluble Fc-tagged ACE2 and the RBD core binding mAb CR3022 were taken along as controls. The V367F substitution and the alanine substitute at this position only had a minor effect on 47D11 antibody binding (Figure 3E), consistent with data showing that this polymorphism had no effect on neutralisation of SARS2-S pseudo type virus (Figure 3I). Collectively, this indicates that 47D11 would be effective against this SARS-CoV-2 variant. In contrast, all other amino acid substitutions in the hydrophobic core not only reduced cell-surface binding by 47D11 (Figure 3E), but also prevented binding of ACE2 and the core targeting antibody CR3022, despite being distal to their respective interaction sites (Figure 3F-G and Figure S4A-B). Total cellular expression of mutants was comparable to wildtype spike protein as demonstrated by an antibody targeting the C-terminal appended Flag-tag on the spike proteins (Figure 3H), suggesting that mutations in the RBD hydrophobic core have a detrimental effect on protein folding, compromising the tertiary structure of the RBD. A recent study reported deep mutational scanning of SARS2-S RBD residues, revealing how mutation of each of the RBD residues affects expression of folded protein and its affinity for ACE2(*46*). When the mean mutation effect on expression was mapped on the 47D11 bound RBD, we observed that the hydrophobic pocket, targeted by 47D11, is highly mutationally constrained (Figure 3J). Another SARS-CoV and SARS-CoV-2 neutralising antibody, S309, targets a similar region to 47D11, but here the orientation of the N343 glycan prohibits access to the hydrophobic pocket, similarly to apo structures (Figure 3K)(*35*). The 47D11 epitope is distinct from other reported RBD-core targeting antibodies/nanobodies, such as CR3022, H014 and VHH-72 (Figure S4)(*27, 38, 47*).

Comparative sequence analysis revealed that the 47D11 epitope is highly conserved across circulating SARS-like viruses (Figure 4A and S5). This is in contrast to the ACE2 binding region which exhibits the greatest sequence variability. In order to assess whether 47D11 has broad reactivity, we recombinantly expressed the RBD from WIV16, HKU3-3 and HKU9-3 and analysed 47D11 binding to these related sarbecoviruses. The results demonstrated that 47D11 can bind to the WIV16 RBD with similar affinity to SARS-S and SARS2-S (Figure 4B). Closer inspections of the aligned RBD amino acid sequences revealed that the N343 glycosylation site, as well as the hydrophobic pocket, are strictly conserved between these three SARS-like viruses (Figure 4C). Therefore, the lack of binding observed for HKU3-3 and HKU9-3 may be due to epitope adjacent sequence differences which preclude binding of the CDRH3 loop. Nevertheless, the potent binding observed for WIV16 underscores the potential of 47D11 as a treatment for future outbreaks caused by SARS-like viruses.

**Figure 4:**
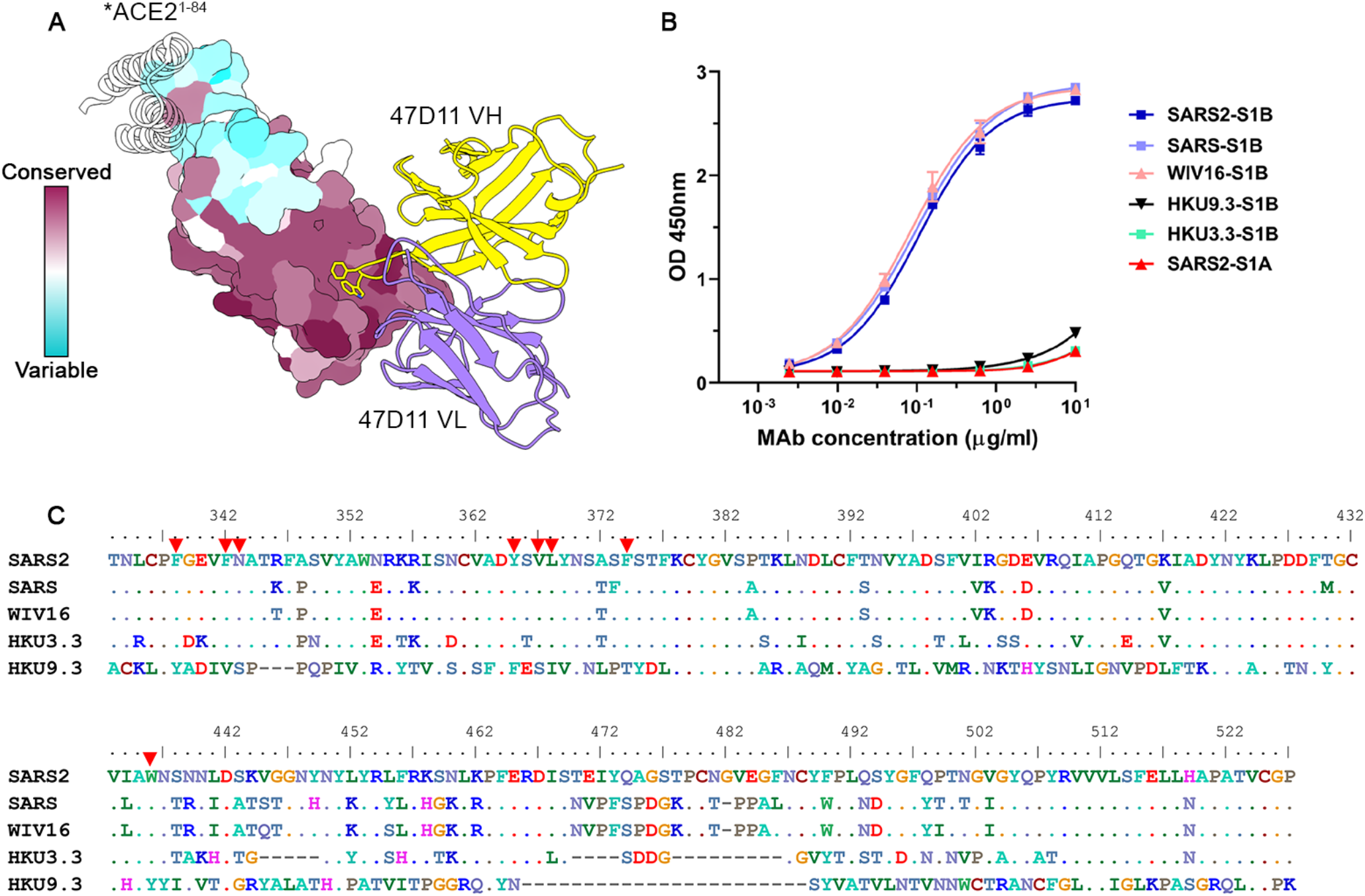
The 47D11 epitope is conserved in SARS-like viruses. A) Surface representation of the 47D11 bound RBD coloured according to sequence conservation across SARS-1, SARS2 and 11 SARS-like viruses (Supplementary Figure 5). The 47D11 Fab variable chains are shown as a ribbon diagram and coloured grey. Heavy chain residues W102 and F103 are shown as sticks. For comparison, residues 1-84 of the RBD bound ACE2 (PDB ID: 6M0J) are shown as a silhouette. B) ELISA-binding curves of 47D11 to the S1B domain of SARS, SARS2, WIV16, HKU3-3 and HKU9-3. The average ±SD from two independent experiments with technical duplicates is shown. C) Aligned RBD sequences of SARS2, SARS-1, WIV16, HKU3-3 and HKU9-3. Key residues in the 47D11 epitope are indicated by red arrows.

In conclusion, our structural and functional analyses demonstrate that 47D11 is able to unmask a conserved and mutationally constrained epitope on the SARS-CoV-2 RBD by stabilising the N343 glycan in an upright conformation. Once this site is exposed, the CDRH3 loop is able to insert two aromatic residues into the hydrophobic core of the RBD, inducing conformational changes which lead to the formation of a 55 Å^3^ cavity. This cryptic site offers an attractive target for design of vaccines and targeted therapeutics. The 47D11 epitope is distal to the ACE2 receptor binding motif, rationalising its ability to cross-neutralize SARS-CoV and SARS-CoV-2 independently of receptor-binding inhibition. Our structural analysis also shows that 47D11 exhibits differing conformational selectivity for the SARS-S and SARS2-S, providing a possible explanation for the differences in observed binding affinity. Two recently described SARS-CoV-2 specific mAbs, C144 and S2M11, recognise quaternary epitopes which partially overlap with 47D11, and lock the SARS-CoV-2 spike in the fully closed conformation(48, 49). In contrast, 47D11 selects for the partially open conformation of the SARS-CoV-2 spike protein, suggesting that it may render the spike more susceptible to other monoclonal antibodies which target the exposed receptor-binding subdomain, making it a prime candidate for combination treatment. Antibody combinations targeting non-overlapping epitopes may act synergistically permitting a lower dosage and an increased barrier to immune escape(*49*). Genetic diversity for SARS2 is currently limited, as the virus has gone through a genetic bottleneck during the singular animal-to-human spill-over event. However, the genetic/antigenic variation will increase in time, as observed for the endemic human coronavirus HCoV-229E, which exhibits cumulative sequence variation in the RBD loops which engage its cellular receptor(50). The seemingly limited mutational space of the 47D11 epitope, in addition to its cross-reactivity within the *Sarbecovirus* subgenus, may confer the antibody sustainable applicability in neutralizing a wide range of future-emerging virus variants.

## Methods

### Expression and purification of coronavirus spike proteins

To express the prefusion spike ectodomain, gene encoding residues 1–1200 of SARS2 S (GenBank: QHD43416.1) with proline substitutions at residues 986 and 987, a “AAARS” substitution at the furin cleavage site (residues 682–685) and residues 1-1160 of SARS S (GenBank: AAP13567.1) with proline substitutions at residues 956 and 957, a C-terminal T4 fibritin trimerization motif, a StrepTag was synthesized and cloned into the mammalian expression vector pCAGGS. Similarly, pCAGGS expression vectors encoding S1 or its subdomain S1_B_ of SARS (S1, residues 1-676; S1_B_, residues, 325-533), and SARS2 (S1, residues 1-682; S1_B_, residues 333-527) C-terminally tagged with Fc domain of human or mouse IgG or Strep-tag were generated as described before(*41*). Recombinant proteins and antibody 47D11 were expressed transiently in FreeStyle™ 293-F Cells (Thermo Fisher Scientific) and affinity purified from the culture supernatant by protein-A sepharose beads (GE Healthcare) or streptactin beads (IBA) purification. Purity and integrity of all purified recombinant proteins was checked by coomassie stained SDS-PAGE.

### Pseudotyped virus neutralization assay

Neutralization with SARS2-S VSV pseudotyped viruses was performed as described previously (41). HEK-293T cells were transfected with pCAGGS expression vectors encoding SARS2-S carrying a 18-a.a. cytoplasmic tail truncation, respectively. One day post transfection, cells were infected with the VSV-G pseudotyped VSVΔG expressing the firefly (*Photinus pyralis*) luciferase. Twenty-four hours later, cell supernatants containing SARS2-S pseudotyped VSV particles were harvested and titrated on African green monkey kidney VeroE6 (ATCC#CRL-1586) cells. In the virus neutralization assay, mAbs were threefold serially diluted and mixed with an equal volume of pseudotyped VSV particles and incubated for 1 hour at room temperature (RT). The virus/antibody mix was subsequently added to confluent VeroE6 monolayers in 96-well plate, and incubated at 37°C. After 24 hours, cells were washed and lysis buffer (Promega) was added. Luciferase activity was measured on a Berthold Centro LB 960 plate luminometer using D-luciferin as a substrate (Promega). The percentage of infectivity was calculated as ratio of luciferase readout in the presence of mAbs normalized to luciferase readout in the absence of mAb. The half maximal inhibitory concentration (IC_50_) was determined using 4-parameter logistic regression (GraphPad Prism version 8).

### ELISA analysis of antibody binding to CoV spike antigens

ELISA was performed as described previously (41). Briefly, NUNC Maxisorp plates (Thermo Scientific) coated with equimolar antigen amounts were blocked with 3% bovine serum albumin (Bio-Connect) in PBS containing 0.1% Tween-20 at RT for 2 hours. Fourfold serial dilutions of mAbs starting at 10 μg/ml (diluted in blocking buffer) were added and plates were incubated for 1 hour at RT. Plates were washed three times and incubated with horseradish peroxidase (HRP)-conjugated goat anti-human secondary antibody (ITK Southern Biotech) diluted 1:2000 in blocking buffer for 1 hour at RT. An HRP-conjugated anti-StrepMAb (IBA) antibody was used to corroborate equimolar coating of the Strep-tagged spike antigens. HRP activity was measured at 450 nanometer using tetramethylbenzidine substrate (BioFX) using an ELISA plate reader (EL-808, Biotek). Half-maximum effective concentration (EC_50_) binding values were calculated by non-linear regression analysis on the binding curves using GraphPad Prism (version 8).

### Preparation of Fab-47D11 from IgG

47D11 Fab was digested from IgG with papain using a Pierce Fab Preparation Kit (Thermo Fisher Scientific), following the manufacturer’s standard protocol.

### Cryo-EM sample preparation and data collection

3 μL of SARS2-S or SARS-S at 1.6 mg/mL was mixed with 0.85 μL of Fab 47D11 at 4 mg/mL and incubated for 50 s at RT. The sample was applied onto a freshly glow discharged R1.2/1.3 Quantifoil grid in a Vitrobot Mark IV (Thermo Fisher Scientific) chamber pre-equilibrated at 4°C and 100% humidity. The grid was immediately blotted at force 0 for 5 s and plunged into liquid ethane. Data was acquired on a 200 kV Talos Arctica (Thermo Fisher Scientific) equipped with a Gatan K2 Summit direct detector and Gatan Quantum energy filter operated in zero-loss mode with a 20 eV slit width. To account for the preferred orientation exhibited by the spike ectodomains, automated data collection at tilts 0°, 20°, and 30° was carried out using EPU 2 software (Thermo Fisher Scientific), and data at tilt 40° using SerialEM(51). A nominal magnification of 130,000x, corresponding to an effective pixel size of 1.08 Å, was used. Movies were acquired in counting mode with a total dose of 40e/Å^2^ distributed over 50 frames. 4,231 movies were acquired for SARS2 and 3,247 movies for SARS-S, with defocus ranging between 0.5 μm and 3 μm.

### Cryo-EM data processing

Single-particle analysis was performed in Relion version 3.1.(*52*). The data was processed in four separate batches, corresponding to the stage tilt angle used for the acquisition. Drift and gain correction were performed with MotionCor2(*53*), CTF parameters were estimated using CTFFind4(*54*) and particles were picked using the Laplacian picker in Relion(*52*). One round of 2D classification was performed on each batch of data and particles belonging to well defined classes were retained. Subsequently, 3D classification was performed, using a 50 Å low-pass filtered partially open conformation as an initial model (EMD-21457,(12)), without imposing symmetry. All particles belonging to Fab bound class were then selected for 3D auto-refinement. Before merging the different batches, iterative rounds of per particle CTF refinement, 3D auto-refinement and post-processing were used to account for the stage tilt used during data collection. The refined particle star files from each batch were then combined and subjected to a final round of 3D auto refinement, per particle defocus estimation, 3D auto-refinement and post processing, both with and without imposed C3 symmetry. Overviews of the single-particle image processing pipelines are shown in supplementary figure 6 and 7.

### Model building and refinement

UCSF Chimera (version 1.12.0) and Coot (version, 1.0.) were used for model building and analysis(*55, 56*). The SARS2-S model, in the partially open conformation (one RBD up, pdb 6VYB)(*12*), was used for the spike and fitted into our density using the UCSF Chimera ‘Fit in map’ tool(*55*). For SARS a closed protomer of the pdb 6NB6 was used as starting model(*57*). To build a model for the Fab the sequence of the variable regions of the HC and the LC were separately blasted against the pdb. For the HC variable region, the corresponding region of the pdb 6IEB (human monoclonal antibody R15 against RVFV Gn) was used(*58*). The LC variable region was modelled using the pdb 6FG1 as template (Fab Natalizumab)(*59*). For both chains, the query sequence of 47D11 was aligned to the template sequence. Sequence identity was particularly high (87% and 97% for the HC and LC, respectively). Phenix sculptor was used to create an initial model for the Fab chains(*60*), removing the non-aligning regions (notably the CDRH3). This model was fitted into our density and the missing regions were built manually in the density map using Coot(*56*). Models were refined against the respective EM density maps using Phenix Real Space Refinement and Isolde(*61*)(*62*), and validated with MolProbity and Privateer (glycans)(*63–65*).

### Analysis and visualisation

PDBePISA was used to identify spike residues interacting with 47D11(*66*). Surface colouring of the SARS-CoV-2 RBD using the Kyte-Doolittle hydrophobicity scale was performed in UCSF chimera(*55*). Volume measurements were performed using CASTp 3.0, using a probe radius of 1.2 Å(*67*). In order to colour the 47D11 bound RBD surface according to each residues mean mutational effect on expression, the pdb file was populated with the mean mutation effect on expression values described by Starr et al(*46*). The UCSF Chimera ‘MatchMaker’ tool was used to obtain RMSD values, using default settings. Figures were generated using UCSF Chimera(*55*) and UCSF ChimeraX(*68*).

## Data availability

Coordinates for the 47D11-bound SARS-CoV and SARS-CoV-2 spike proteins are deposited in the Protein Data Bank under accession codes 7AKJ and 7AKD, respectively. The corresponding EM density maps have been deposited to the Electron Microscopy Data Bank under the accessions EMD-11813 and EMD-11812. All reagents and relevant data are available from the authors upon request.

## Acknowledgements

This work was supported by the European Research Council under the European Union’s Horizon2020 Programme (ERC Consolidator Grant Agreement 724425 - BENDER). J.F. and D.L.H. are funded from the European Union’s Horizon 2020 research and innovation program under the Marie Skłodowska-Curie grant agreement (No 792575 and 842333). J.F. and D.L.H. also hold EMBO non-stipendiary long-term Fellowships (ALTF 948-2017 and ALTF 1172-2018). Research reported in this publication was supported by a China Scholarship Council grant to C.W. (CSC201708620178). We thank Dr Mihajlo Vanevic for IT support.

## Author contributions

J.F., D.L.H., C.W., W.L., F.J.M.K., F.F. and B.J.B. conceived, designed, and coordinated the study. J.F., D.L.H., C.W., W.L., I.D., G.O., and S.H. conducted the experiments. F.J.M.K, F.F. and B.J.B. supervised part of the experiments. All authors contributed to the interpretations and conclusions presented. J.F. D.L.H. and B.J.B. wrote the manuscript, J.F., D.L.H, C.W., W.L. F.F. and B.J.B participated in editing the manuscript.

## Corresponding authors

Correspondence to Friedrich Förster and Berend-Jan Bosch.

