## Supplementary Information for "A human monoclonal antibody targeting a conserved pocket in the SARS-CoV-2 receptor-binding domain core"

**Fedry et al.**

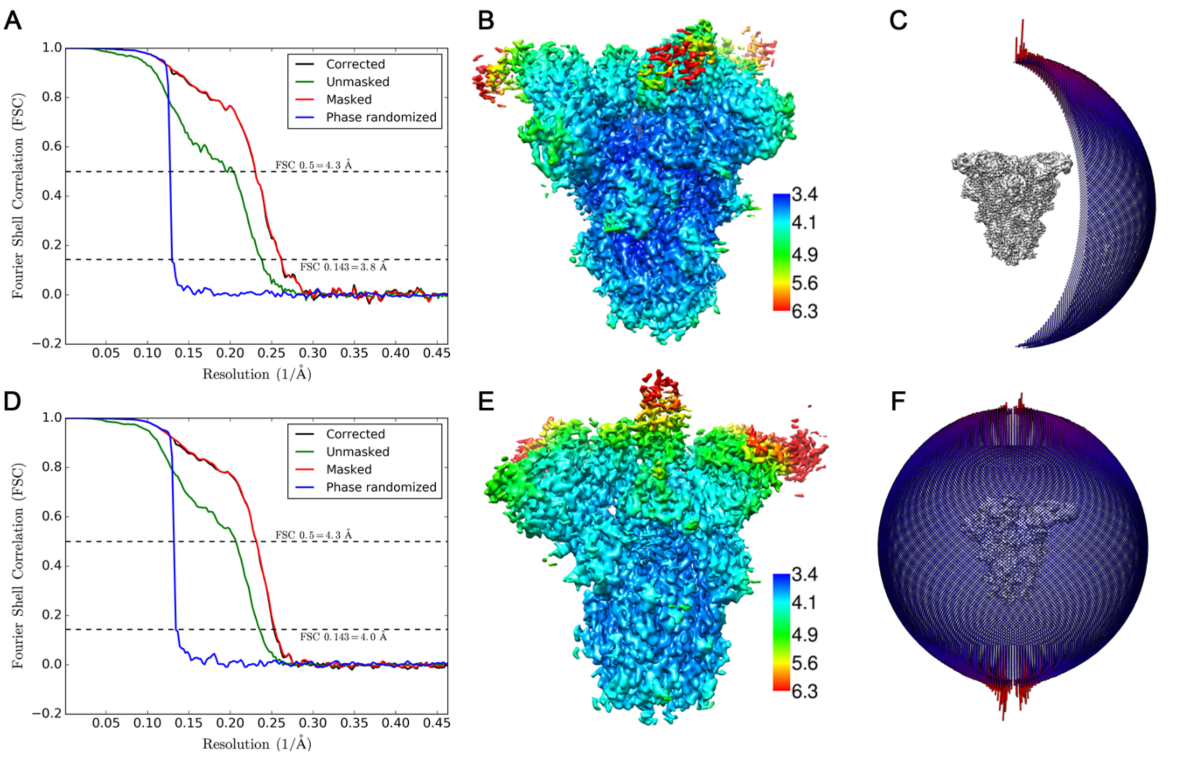

**Figure S1:** A) Gold-standard Fourier shell correlation (FSC) curve generated from the independent half maps contributing to the 3.8 Å global resolution density map of the SARS-CoV spike in complex with the 47D11 antibody Fab fragment. B) Local resolution filtered EM density map for the C3 refined SARS-CoV spike:47D11 complex, coloured according to local resolution which was calculated in Relion3.1. C) Angular distribution plot of the final C3 refined SARS-CoV spike EM density map. D) Gold-standard FSC curve generated from the independent half maps contributing to the 4 Å global resolution density map of the SARS-CoV-2 spike in complex with the 47D11 antibody Fab fragment. E) Local resolution filtered EM density map for the C1 refined SARS-CoV-2 spike:47D11 complex, coloured according to local resolution which was calculated in Relion3.1. F) Angular distribution plot of the final C1 refined SARS-CoV-2 spike EM density map.

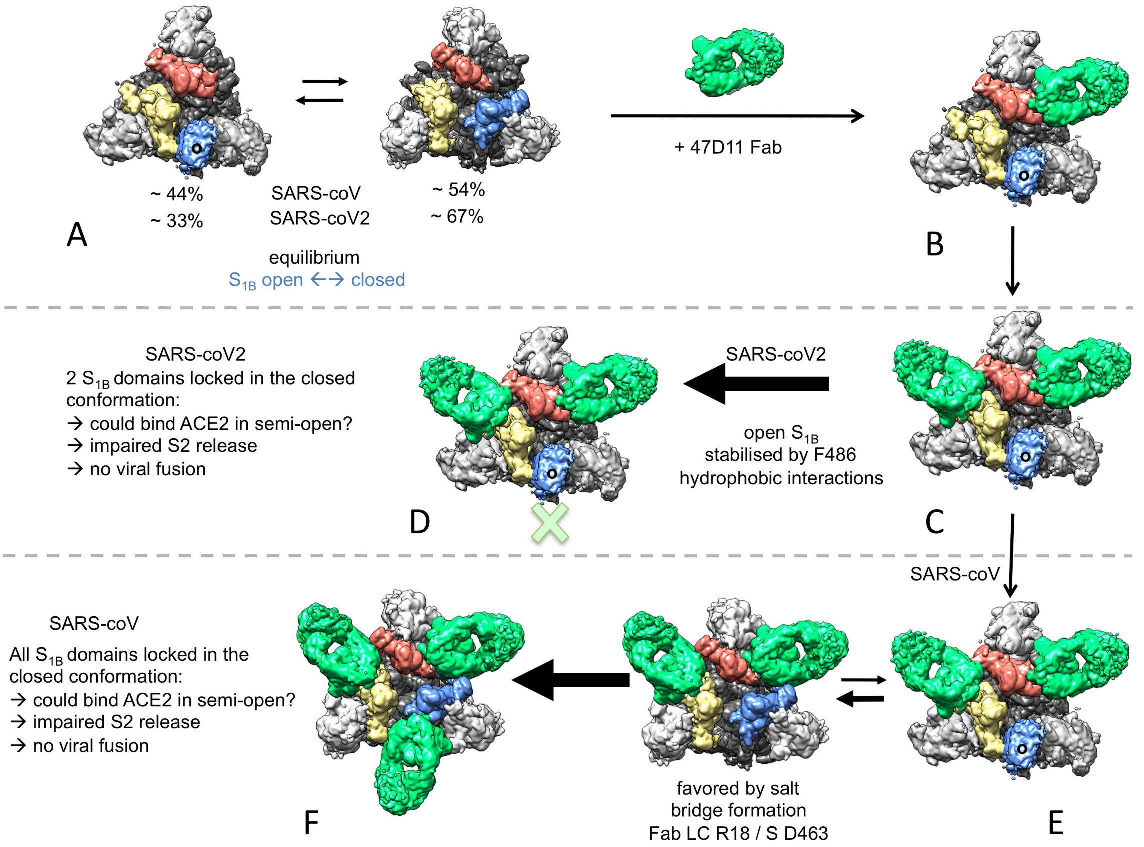

**Figure S2: Proposed mechanism of 47D11 binding to SARS-CoV and SARS-CoV-2**

A) The Spike protein in solution exists in equilibrium between the closed and the partially open conformation. The $S_{1A}$ (NTD) domains are colored in light gray, the $S_{1B}$ domains are in salmon, beige and blue and the rest of the spike protein is in dark gray. For clarity, the letter « o » is added on the blue $S_{1B}$ domain when in the open conformation. B) The 47D11 Fab is added and binding of a first Fab on the salmon closed $S_{1B}$, right next to the open blue S1B is favored (47D11 epitope is more accessible). C) A second 47D11 Fab binds the second closed beige $S_{1B}$ domain. In the case of SARS-CoV it is favored by the formation of an extra salt bridge between this second Fab and the previous salmon $S_{1B}$ distal loop (Fab LC R18 / S D463). D) For SARS-CoV-2 the blue $S_{1B}$ is stabilized in the open conformation, likely by hydrophobic interactions formed by the $F_{486}$ of the distal loop with the open $S_{1B}$ and possible clashes of its RBR loop with the first bound Fab upon closing. No 47D11 Fab can bind to this open third $S_{1B}$ as this would result in severe clashes with the neighbouring $S_{1A}$. This is the final conformation stabilized by 47D11 for SARS-CoV-2 S. E) for SARS-CoV S, an equilibrium can exist in the form with 2 bound 47D11 Fabs, where the blue $S_{1B}$ domain could open and close. The closed form is stabilised by the formation of the extra salt bridge. F) When the blue $S_{1B}$ domain is closed, a third 47D11 Fab can bind on it. This is the final conformation stabilised by the 47D11 Fab for SARS-CoV. In both SARS-CoV and SARS-CoV-2 the 47D11 Fab prevents the possibility of fully opening the 3 $S_{1B}$ domains (as 2 or 3 of them are bound by 47D11 and cannot fully open anymore because of severe clashes with the next $S_{1A}$). They may be able to bind ACE2 in a partially open conformation that would impair the timely release of the fusogenic $S_2$ domain and thereby prevent viral fusion.

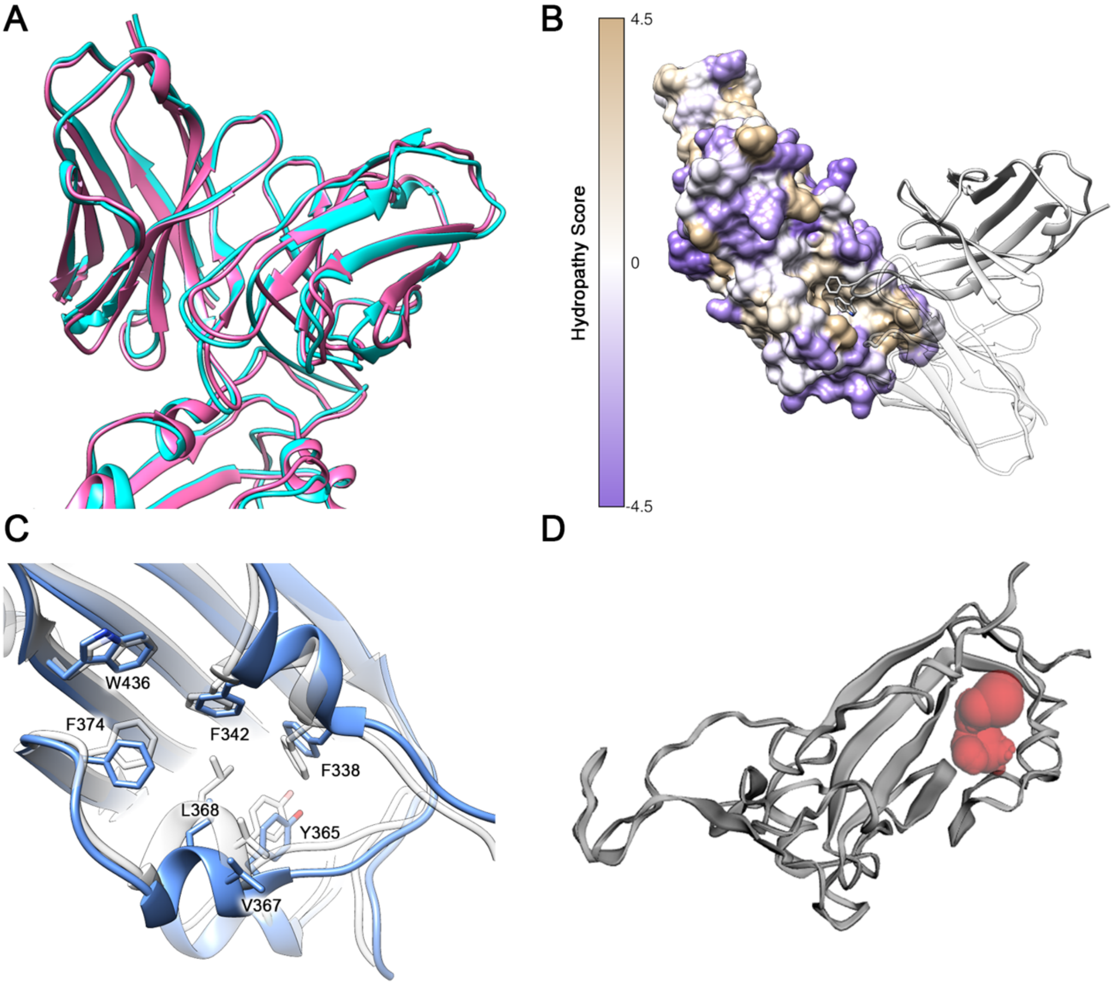

**Figure S3**: A) Superposed atomic coordinates for the 47D11 bound SARS-S and SARS2-S RBD, coloured cyan and pink, respectively. B) The SARS2-S RBD shown as a surface representation and coloured according to the Kyte-Doolittle scale, where the most hydrophobic residues are coloured tan and the most hydrophilic residues are coloured purple. The 47D11 Fab fragment is shown as a ribbon diagram, with the light chain shown semi-transparent. C) Zoomed in view of the hydrophobic pocket for the 47D11 bound SARS2 RBD (blue) overlaid with the equivalent region from an apo structure (PDB ID: 6VYB). D) Ribbon diagram of the 47D11 bound RBD with the 55 Å$^3$ solvent accessible cavity, generated using CASTp 3.0, shown as red spheres(*1*).

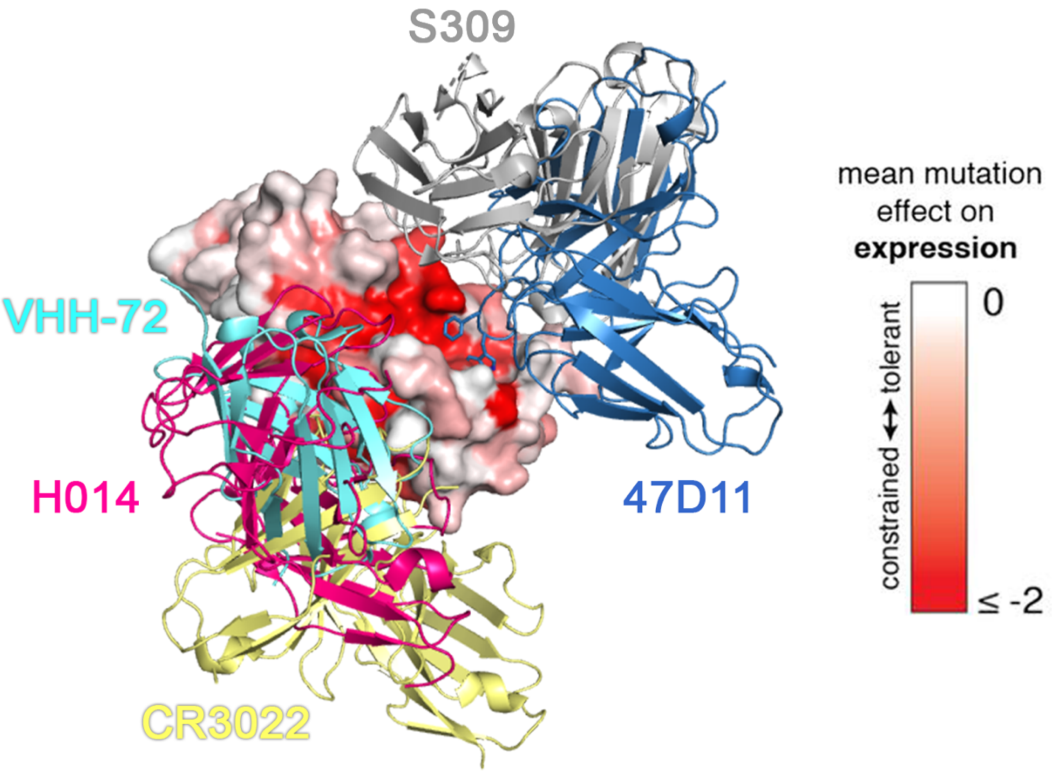

**Figure S4:** Surface rendering of the SARS2-S RBD, coloured according to the mean mutation effect on expression (red indicates more constrained)(*2*), with binding positions of 47D11, S309 (PDB ID: 6WPS)(*3*), H104 (PDB ID: 7CAH)(*4*), CR3022 (PDB ID: 6W41)(*5*) and VHH-72 (PDB ID: 6WAQ)(*6*) overlaid.

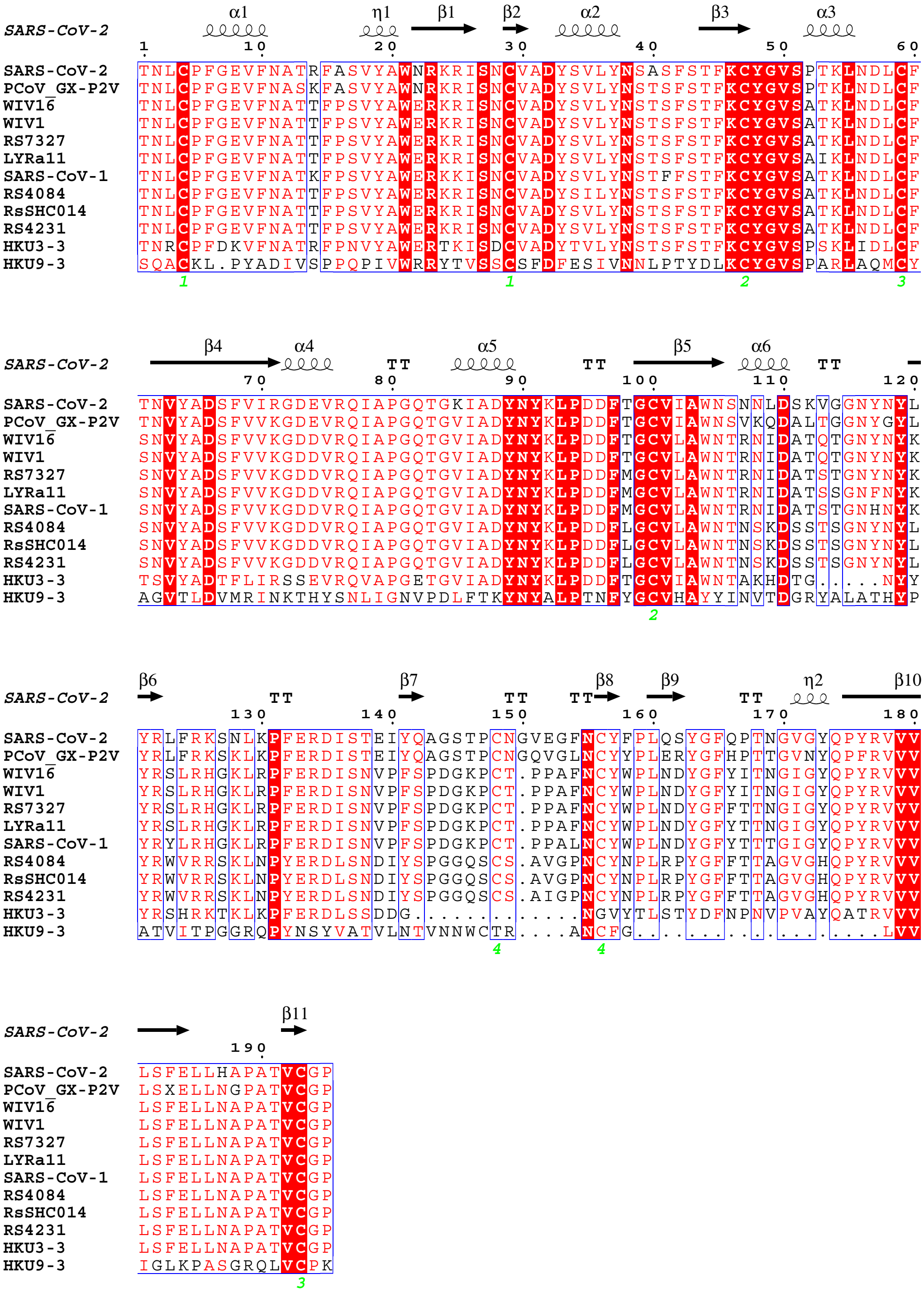

**Figure S5:** Multiple sequence alignment of the RBD residues from SARS-CoV, SARS-CoV-2 and 11 SARS-like viruses. The sequence alignment was performed using Clustal Omega(*7*) and the image was generated by ESPript 3.01(*8*). The secondary structure assignment, based on the SARS2 RBD, is shown.

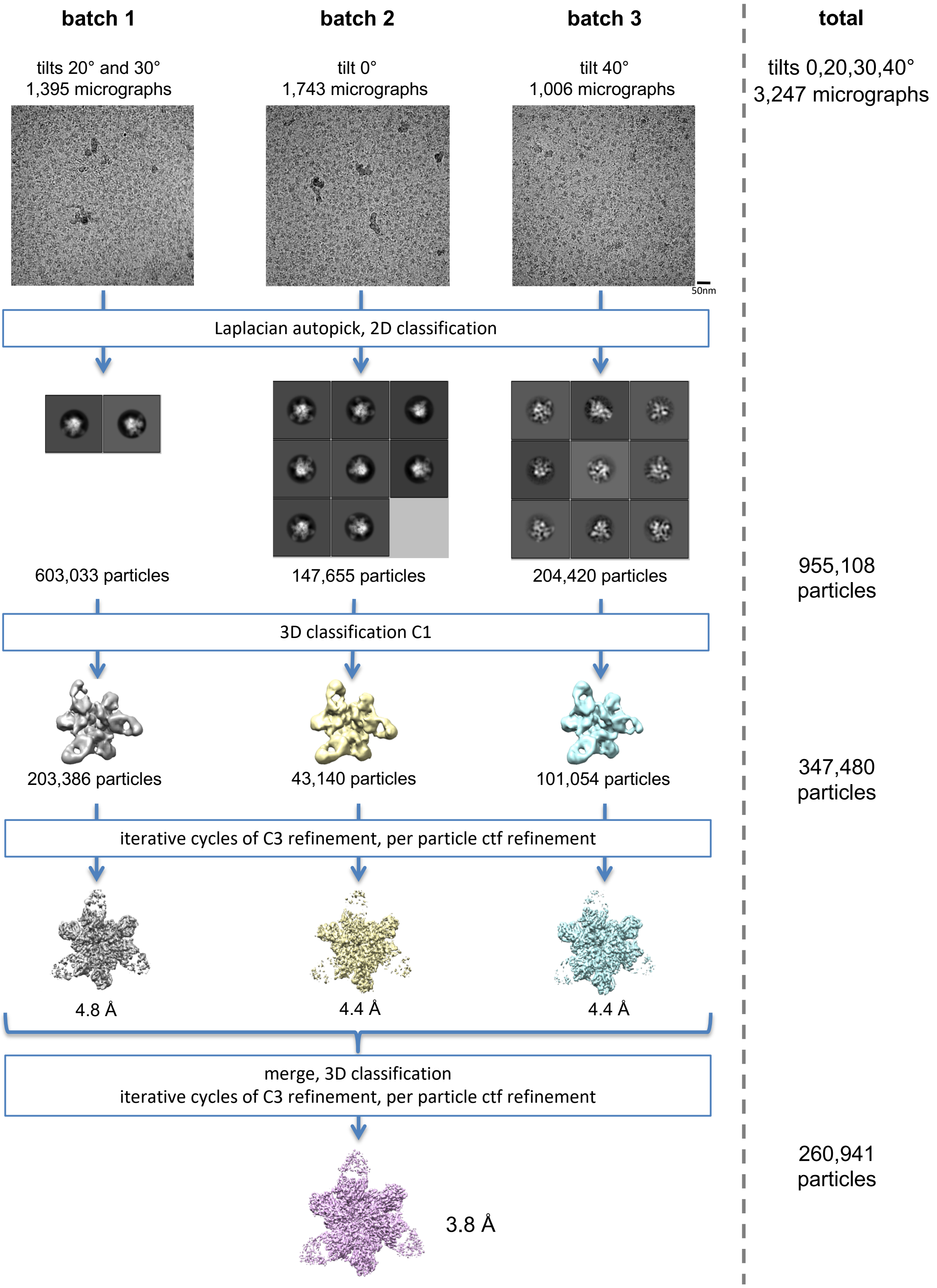

**Figure S6:** Single-particle cryo-EM image processing workflow for the SARS-CoV:47D11 complex.

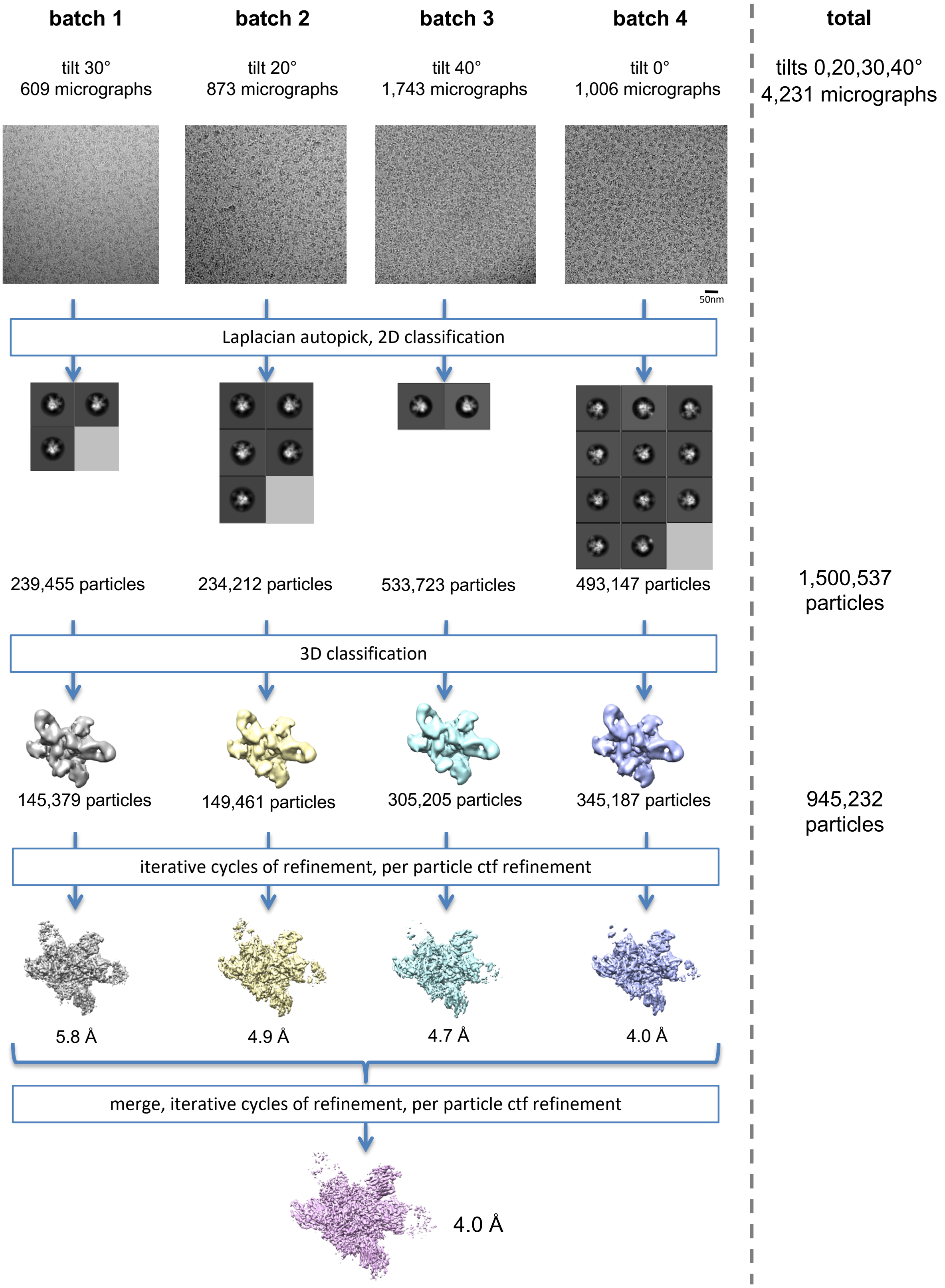

**Figure S7:** Single-particle cryo-EM image processing workflow for the SARS-CoV-2:47D11 complex.

**Table S1:** Data collection, image processing and refinement information.

| | SARS-coV + 47D11 | SARS-coV2 + 47D11 |
|---|---|---|
| **Data Collection and processing** | | |
| Magnification | 130 000 | 130 000 |
| Voltage (kV) | 200 | 200 |
| Electron exposure (e-/Å$^2$) | 40 | 40 |
| Defocus range (μm) | 0.5-2.5 | 0.5-2.5 |
| Pixel size (Å) | 1.08 | 1.08 |
| Symmetry imposed | C3 | C1 |
| Initial particle images (no.) | 955 108 | 1 500 537 |
| Final particle images (no.) | 260 941 | 945 232 |
| Map resolution (Å) | 3.8 | 4.0 |
| FSC threshold | 0.143 | 0.143 |
| Map resolution range (Å) | 3.4-6.3 | 3.4-6.3 |
| **Refinement** | | |
| Initial model used (PDB code) | 6NB6 | 6VYB |
| Model resolution (Å) | 3.9 | 4.0 |
| FSC threshold | 0.5 | 0.5 |
| Map sharpening B factor (Å$^2$) | -168 | -154 |
| Model composition | | |
| Amino acids | 3930 | 3404 |
| Glycans | 96 | 79 |
| mean B factors (Å$^2$) | | |
| Protein | 42.4 | 44.1 |
| Glycans | 91.4 | 83.9 |
| R.m.s deviations | | |
| Bond lengths (Å) | 0.005 | 0.003 |
| Bond angles (°) | 0.826 | 0.624 |
| Validation | | |
| Molprobity score | 1.49 | 1.21 |
| Clashscore | 5.99 | 2.90 |
| Poor rotamers (%) | 0.09 | 0.21 |
| Ramachandran plot | | |
| Favored (%) | 97.12 | 97.31 |
| Allowed (%) | 2.88 | 2.69 |
| Disallowed (%) | 0 | 0 |
